# Skeletal remains of a freshwater jalodontid chondrichthyan from the Upper Triassic Arden Sandstone Formation of Warwickshire, UK

**DOI:** 10.64898/2026.09.08.749451

**Authors:** Ryan R. Tokeley, Richard P. Dearden, Stuart D. Burley, Jonathan D Radley, Ivan Sansom

**Affiliations:** University of Birmingham, Birmingham, UK; Naturalis Biodiversity Center, Leiden, The Netherlands; Basin Dynamics Research Group, Keele University, Keele, UK

## Abstract

The Triassic was a significant period in the evolution of modern chondrichthyans (sharks, rays and chimaeras), bridging the gap between Paleozoic chondrichthyans and more recognizably modern Mesozoic forms. Only a handful of Triassic skeletal fossils of chondrichthyans are known worldwide, their calcified cartilages provide rare insights into the anatomy, evolution, and lifestyle of these important taxa. Here we use computed tomographic methods to describe six chondrichthyan skeletal fossils from the Carnian (Upper Triassic) Arden Sandstone Formation of Shrewley, Warwickshire, UK, consisting of a partial braincase, jaws, pectoral fin radials, caudal fin hypurals, and dorsal fin basal cartilages, and characterize their paleoenvironment. We assign the braincase and jaws, and tentatively the post-cranial remains, to the jalodontid taxon *Keuperodus brodiei*, previously known only from teeth. This represents the first interpretable cranial cartilages of the Order Jalodontiformes as well as the first known occurrence of articulated chondrichthyan skeletal remains from the Carnian. The partial braincase of *Keuperodus brodiei* preserves an unusual basicranial circulation, almost completely embedded in the basicranium, and the jaws have an elongate extended shape; both features are akin to Paleozoic phoebodontid chondrichthyans. Phylogenetic analysis recovers *Keuperodus brodiei* as a stem-group elasmobranch with a close relationship to phoebodonts, showing jalodontids to be one of several stem-group elasmobranch lineages to have independently survived the end-Permian mass extinction. However, unlike Phoebodontiformes and all other Jalodontiformes, *Keuperodus brodiei* occupied freshwater lake shorelines and associated fluvial feeder channels, revealing a possible freshwater refugium for Jalodontiformes towards the end of their stratigraphic range.

## Introduction

The Triassic is a period of major significance for the early evolution of modern elasmobranchs, spanning important early divergences between extant cartilaginous fish groups (Marion et al. 2024; Sorenson et al., 2014; Stein et al., 2018). The Triassic also bridges the gap between the late Paleozoic, where advances in x-ray based imaging methods have formed an increasingly sophisticated picture of early crown-group elasmobranchs (Bronson et al., 2018; Coates & Tietjen, 2017; Frey et al., 2019; Pradel et al., 2014), and the mid-late Mesozoic, where articulated elasmobranchs from fossil-lagerstätten shed light on the origins of living elasmobranch Orders (Dearden et al., 2025; Jambura et al., 2023; Stumpf et al. 2026; Vullo et al., 2024). As a result, Triassic elasmobranchs have the potential to inform our understanding of the transition from Palaeozoic to modern elasmobranch groups as well as that of their recovery from the end-Permian mass extinction (EPME) and the effect this had on their evolutionary history. Isolated teeth, fin spines and dermal remains demonstrate that Triassic chondrichthyan faunas included members of groups which went on to flourish later in the Mesozoic such as hybodonts (Prasad et al., 2008; Wen et al., 2023), cryptic taxa with unclear relationships to extant elasmobranchs (Cappetta, 1987; Sykes, 1975) as well as remnants of Paleozoic groups (Ivanov et al., 2021; Johnson, 1980; Mutter and Rieber, 2005). However, only very few of these Triassic taxa are known from skeletal remains, almost all of which are hybodonts or hybodont-like taxa (Maisey, 2011; Mutter et al., 2007; Mutter and Neuman, 2008; Rieppel, 1982; Romano and Brinkman, 2010; Thomson, 1982). The only Late Triassic endoskeletal remains known are the Rhaetian *Raineria osswaldi*, a rostrum of uncertain taxonomic affinity (Schultz, 2013), and isolated patches of tesselate cartilage from the Upper Triassic of Arizona and Texas suggested to belong to a xenacanthiform or hybodontiform (Velin et al., 2026).

Jalodontiformes are a chondrichthyan Order spanning the Upper Devonian to the Upper Triassic which are almost entirely known from isolated teeth (Ivanov et al., 2021). Previously included within Phoebodontiformes due to their tricuspid tooth morphology, Ivanov et al. (2021) placed jalodontids in a newly erected order of Jalodontiformes based on their previously unrecognized distinctive tooth morphology. Both phoebodontids and jalodontids are known from localities worldwide and overlap in their stratigraphic distribution, with phoebodontids known from the Middle Devonian to the Mississipian (Ginter et al. 2010). The stratigraphically earliest known jalodontid, *Jalodus australiensis*, is known from Famennian marine deposits contemporaneous to those preserving *Phoebodus saidselachus* (Frey et al., 2019; Ivanov et al., 2021; Long, 1990). However, unlike jalodontids, phoebodontids are known from extensive three-dimensionally preserved articulated remains of *Phoebodus saidselachus* (Frey et al., 2019; Klug et al., 2026) as well as flattened specimens of the Mississippian *Thrinacoselache gracia* (Ginter & Turner, 2010; Grogan & Lund, 2008). By contrast, the only articulated jalodontid remains are known from a single specimen of the Permian taxon *Adamantina benedictae* (Bendix-Almgreen, 1993) which preserves scales and teeth but no interpretable endoskeletal remains. As such, the relationships of Jalodontiformes to other Paleozoic shark groups and the significance of their survival into the Triassic remain obscure.

The stratigraphically youngest Jalodontiformes are teeth assigned to *Keuperodus brodiei*, found in the Upper Triassic (Carnian) Arden Sandstone Formation of Shrewley, Warwickshire, UK, and the contemporaneous Grabfeld Formation of Gaildorf-Münster, Germany (Ivanov et al., 2021). The Arden Sandstone Formation, the *Keuperodus brodiei* type locality, has also yielded teeth, dorsal fin spines and cephalic spines attributed to the hybodont *Palaeobates keuperinus* (Burley et al., 2023; Old et al., 1991), but also several undescribed specimens of calcified cartilage representing pieces of chondrichthyan endoskeleton (Brodie, 1887, 1893). In this study, we use computed tomography (µCT) to describe six of these calcified cartilage specimens, representing the remains of one or more chondrichthyan individuals collected from the Arden Sandstone of Shrewley during the late nineteenth century. Two of these specimens are confidently assigned to *Keuperodus brodiei* and the other four we assign to Elasmobranchii *incertae sedis*.

## Materials and Methods

### Museum Collections and Abbreviations

NHMUK, Natural History Museum, London; SMNS, Staatliches Museum für Naturkunde Stuttgart, Germany; WARMS, Warwickshire Museum, Warwick, England.

### Geological and Historical Setting

The Arden Sandstone Formation is assigned to the upper part of the Carnian Stage of the Upper Triassic, deposited some 234-230 million years ago, seen at outcrop and in boreholes in central and western England, north of the Mendip Hills (Burley et al., 2023; Old et al., 1991). The unit is a distinctive, pale-green colored, arenaceous deposit marking a temporary divergence from the red mudstones characteristic of the Mercia Mudstone Group. The Arden Sandstone Formation thus separates the lower Sidmouth Mudstone Formation from the overlying Branscombe Mudstone Formation (Burley et al., 2023; Newell, 2024); the over and underlying formations reflecting deposition on continental dryland desert floodplains, with soil profiles (Milroy et al., 2019). The distinctive sedimentological signature of the Arden Sandstone reflects expansion of a shallow, oxygenated, freshwater lake termed ‘Lake Arden’. The lake was large, extending from the Mendips in the south to the Stafford Basin in the north, a distance of ∼100km (Burley et al., 2023) and was fed by small, ephemeral fluvial channels sourced from the flanking residual uplands and crossing a low-relief, desert plain topography. Dissolution breccias, desiccation cracks and the presence of halite pseudomorphs and gypsum nodules indicate that at times Lake Arden was of limited extent and surrounded by evaporitic playa muds (Burley et al., 2023). Lacustrine and fluvial environments attracted a range of fauna and flora whilst promoting fossil preservation, resulting in a relatively diverse and distinct fossil record known from the formation. Body fossils include the previously mentioned jalodont and hybodont sharks, scales and partially complete specimens of the osteichthyans *Semionotus* and *Dictyopyge* and teeth of *Ceratodus*, bones of temnospondyl amphibians, tests of the clam shrimp *Euestheria* and poorly preserved molds of bivalves probably belonging to freshwater unionid taxa (Burley et al. 2023). Plant fossils include stems and cones of *Voltzia*, *Neocalamites*, *Schizoneura* as well as several seed morphotypes all assigned to *Carpolithus*. The ichnofauna is restricted being dominated by small burrows attributed to *Planolites* as well as rare footprint casts of *Chirotherium* and *Rhynchosauroides* (Brodie, 1856, 1870, 1893, 1896; Burley et al., 2023; Matley, 1912; Murchison & Strickland, 1837; Old et al., 1991; Radley, 2006; Wanner, 1921; Warrington & Pollard, 2012). The expansion of Lake Arden broadly correlates with the Carnian Pluvial Episode (CPE), a major climatic shift from arid to humid conditions during the Carnian stage. Although the CPE has been recognized since the 1980’s (Simms & Ruffell 1989, 2018), it has recently gained significance as a driver of major faunal and floral change during the Upper Triassic. (Benton et al., 2018; Bernardi et al., 2018; Dal Corso et al., 2020; Zhang et al., 2023).

The type section of the Arden Sandstone Formation is exposed along the banks of the Grand Union Canal at the village of Shrewley in Warwickshire (SP 21276 67341) (Fig. 1), where the whole formation can be observed, including the contacts with the underlying Sidmouth Mudstone Formation and overlying Branscombe Mudstone Formation. At Shrewley, the base of the Arden Sandstone Formation rests on an eroded surface of the Sidmouth Mudstone Formation. A basal lag of rolled mudstone and dolomite clasts occurs in small erosion hollows formed in the top of the underlying mudstone. The Arden Sandstone Formation is almost 8m thick at Shrewley and coarsens-upwards through rippled silty mudstones and fine-grained sandstones into cross-bedded, medium and locally coarse-grained sandstones with common rip-up mud clasts and quartz and phosphate granules, the latter being rolled phosphatic bone material (Fig. 2). Most of the known fossil specimens from the Arden Sandstone Formation were collected here during the nineteenth century by Reverend Peter Bellinger Brodie (1815 -1897), an English geologist and appointed vicar of the nearby St Laurence’s Church, Rowington. Brodie took a keen interest in the formation and collected a wide range of fossil material from the once active stone quarries along the Grand Union Canal at Shrewley and Rowington (Brodie 1880, 1887, 1893, 1896) which were then described in more detail by Woodward (1889b, 1893).

**FIGURE 1.**
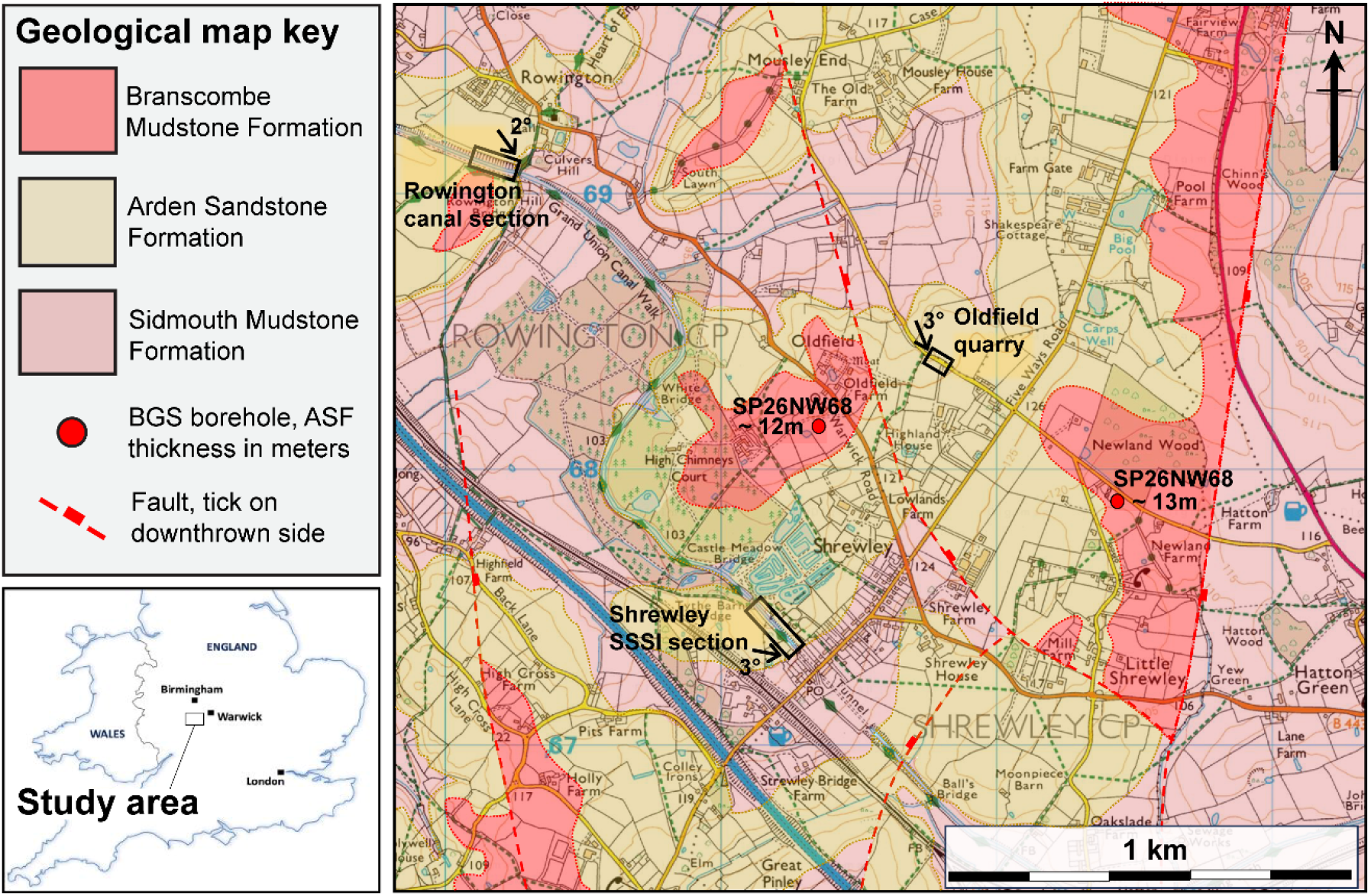
Geological map and stratigraphy of the area around Shrewley, based on the BGS GeoIndex map (https://www.bgs.ac.uk/map-viewers/geoindex-onshore/) modified with the authors own field observations. The geological formations are displayed on the Ordnance Survey Land Ranger map 139. Nearby boreholes report up to 13m of Arden Sandstone Formation being present, but only ∼8m are present in the Shrewley canal section. Regional dip directions are shown for these outcrops. Abbreviations: ASF, Arden Sandstone Formation; BGS: British Geological Survey, SSSI, Site of Special Scientific Interest. (Full page width: 182.033 mm x 111.55 mm)

**FIGURE 2.**
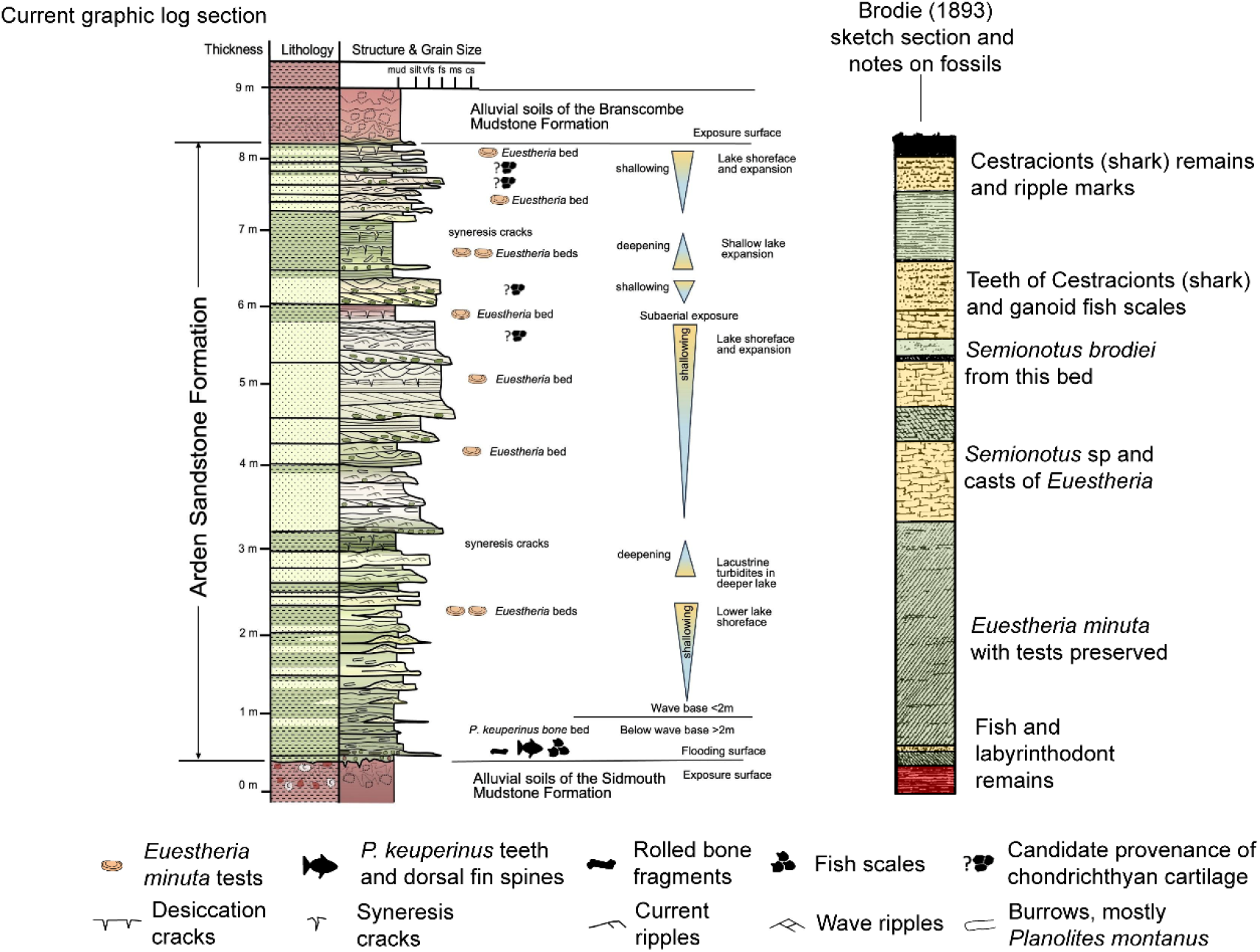
Sedimentary log of the Arden Sandstone Formation at Shrewley showing sedimentary structures and the provenance of fossil remains, constrained with the description made by Brodie in 1893 and his fossil finds. The Arden Sandstone Formation here comprises three, potentially four, shallowing-upward lake sequences and lake deepening events. Many of the chondrichthyan, osteichthyan and amphibian remains occur in the coarse-grained sandstone at the base of the section, recording the initial flooding and establishment of Lake Arden. Candidate beds for the provenance of the calcified cartilage specimens are shown on the log. (Full page width: 182.033 mm x 122 mm)

At the time of writing, the vast majority of Brodie’s collection of Arden Sandstone Formation fossil material is separated between the Warwickshire Museum and the Natural History Museum London with a small number of his specimens located at the British Geological Survey, Lapworth Museum of Geology and Oxford University Museum of Natural History. This historical division of material between Warwick and London includes the specimens referred to *Phoebodus brodiei* (now *Keuperodus brodiei*), and six specimens of undescribed chondrichthyan calcified cartilage, the focus of the current study.

### Fossil Specimens

Six specimens are described here, all of which preserve three-dimensional tessellate calcified cartilage of a yellow-brown color that contrasts with a medium grey, finely arenaceous matrix. Only calcified parts of the skeleton are preserved (Fig. 3). WARMS G136 (Fig. 3a) is held in the Warwickshire Museum collections. The remaining five, NHMUK PV P 7611 and NHMUK PV P 7612a-d (Fig. 3 b-f), are held at the Natural History Museum, London, collections. On being donated WARMS G136 was originally identified as “skin of shark” and in 1981 was relabelled as “Impression of cartilage of skull of *Palaeobates keuperinus*”, presumably on the basis of the abundance of teeth and spines attributed to that taxon from Shrewley. It comprises a single block containing a large, complex piece of cartilage (Fig. 3a). NHMUK PV P 7611 (Fig. 3b) is labelled as “Selachian jaws” and comprises a single block that contains the posterior of a palatoquadrate and Meckel’s cartilage and other isolated fragments of cartilage. NHMUK PV P 7612a (Fig. 3c), PV P 7612b (Fig. 3d), PV P 7612c (Fig. 3e), and PV P 7612d (Fig. 3f) are all recorded as “fragments of cartilage, Selachian” and consist of blocks of fragmentary cartilage material. All of the specimens were collected from Shrewley by Reverend Peter B. Brodie, likely between 1850 and 1887, though the exact dates of collection are not known.

**FIGURE 3.**
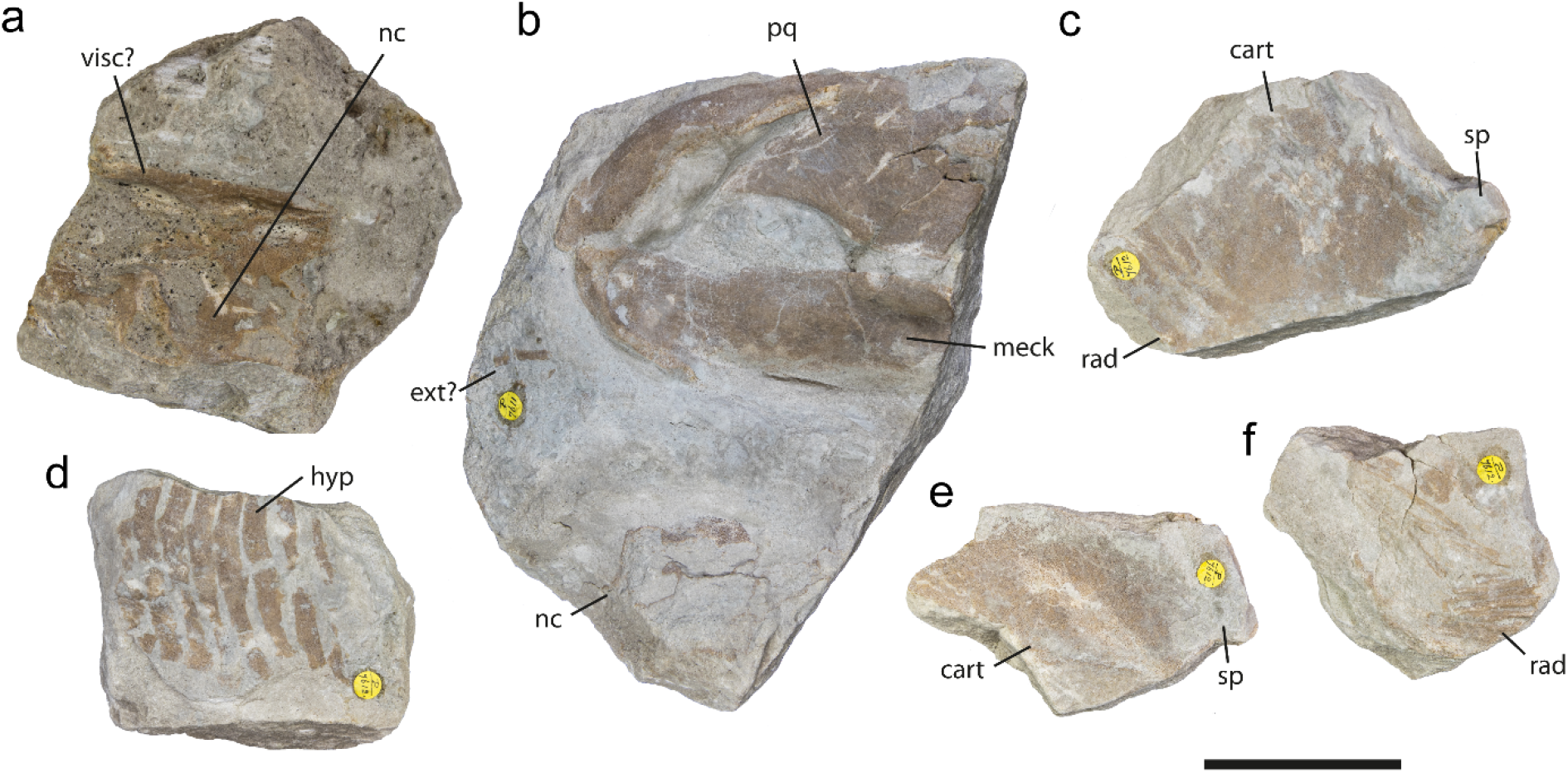
Photographs of chondrichthyan cartilage specimens in this study: a, WARMS G136; b, NHMUK PV P 7611; c, NHMUK PV P 7612a; d, NHMUK PV P 7612b; e, NHMUK PV P 7612c; f, NHMUK PV P 7612d. Scale bar 5 cm. (Full page width: 182.033 mm x 92 mm)

### Specimen Provenance

The cartilage specimens are associated with a fine-to medium-grained, buff-grey colored, cross-bedded sandstone, representing deposits of the shallow lake shoreface. The cross bedding was produced by wind-generated waves that reworked the sediment repeatedly. As a result, the chondrichthyan remains are partially disarticulated, likely having been transported a short distance in a few meters of water depth before being distorted and crushed during burial. Brodie’s (1893) log of the Arden Sandstone section at Shrewley, highlights two sandstone beds containing chondrichthyan (then referred to as “Cestraciont”) fossils in the middle and upper part of the outcrop (Fig. 2). Through field work, we identified four sandstone beds of an identical lithology to the matrix material of all chondrichthyan cartilage specimens (Fig. 2). Although apparently unfossiliferous during the present study (likely due to their current limited outcrop) the sedimentology and stratigraphic position of these beds with respect to Brodie’s “Cestraciont” bearing beds, strongly suggests that these horizons are the most likely source of the chondrichthyan cartilage specimens.

### X-Ray Computed Tomography and Modelling

WARMS G136 (Fig. 3a) was scanned at the University of Birmingham on a Nikon Metrology XT H 225 ST 2x CT scanner. NHMUK PV P 7611 and NHMUK PV P 7612a-d (Fig. 3b-f) were scanned at the Natural History Museum, London, on a Nikon Metrology HMX ST 225 CT scanner. Full details of scanning are given in Supplementary Table 1. A tungsten reflection target was used for all specimens. The scan data was segmented using Materialise Mimics v25.0 via a combination of manual thresholding and interpolation. Threshold ranges between 10,400 and 14,100 GV were used, and separate masks were created for individual skeletal elements. Three-dimensional models were then exported as a wavefront.obj, imported into Blender v4.5 and rendered as high-fidelity models with a three-point lighting setup.

### Phylogenetic Analysis

To investigate the evolutionary relationships of *Keuperodus brodiei* we modified the phylogenetic matrix of Klug et al. (2026). The matrix of Klug et al. (2026) comprised 66 taxa and 230 characters and focuses on chondrichthyans and related taxa with extensively preserved endoskeletons. We modified this matrix by adding one new taxon, *Keuperodus brodiei*, scored on the basis of the interpretation presented here of WARMS G136 and NHMUK PV P 7611 (Fig. 3a,b) as well as Ivanov et al. (2021), i.e. the partial braincase, jaws, and teeth. In addition, we modified scoring for *Phoebodus* character 87, scoring it as having a large otic process present. Finally, we added an additional state for character 116, internal carotid entry “enter from basicranial circuit within the basicranium” which we scored as present in *Phoebodus* and *Keuperodus* based on the interpretation given below. The placoderm genus *Entelognathus* was used as an outgroup taxon. All characters were treated as unordered and of equal weight. Character coding was done via Mesquite v4.02. Phylogenetic analyses used a maximum parsimony approach carried out in TNT v1.6. We ran an initial search of 1000 replicates using RAS and TBR, holding 10 trees from each replicate. We then carried out additional swapping on the resulting trees with TBR, holding a maximum of 20000 trees. We calculated the consistency index (CI) and retention index (RI) for each tree using the stats.run script bundled with TNT. An analysis using a Bayesian approach was carried out in MrBayes v3.2.7, with a likelihood model using gamma rates, variable coding, and 5 beta categories. A variable rate prior was applied. This was run for 15,000,000 generations, with the analysis being ended once the average standard deviation of split frequencies was below 0.01, and stationarity confirmed in Tracer. A 50% burn-in was applied before sampling.

## Results

### Sedimentological setting of the jalodontid chondrichthyan remains

The detailed sedimentological logging of the Arden Sandstone Formation at Shrewley matches remarkably well with the sketch section published by Brodie (1893) and enables the provenance of the jalodontid skeletal remains to be defined with high confidence and the depositional environment to be reconstructed in detail (Figure 2). The Arden Sandstone Formation at Shrewley represents the initial establishment of Lake Arden, with the basal fine sandstones and silty mudstones representing the deepest part of the lake, shallowing into a series of sandy shorelines. The initial 0.5m of silty mudstones were deposited below wave base, but the remainder of the succession was deposited above wave base in the lake as indicated by the abundance of ripples. Lake Arden was large, covering an aerial extent of at least 3000km^2^ (Burley et al., 2023) and had a long fetch which exceeded 100km that would have allowed strong winds and storms to easily create wavelengths of 5-15 meters. Water depths are estimated from ripple wavelength (depth = wavelength/2), suggesting depths in the range 2.5 to 7.5m, and providing a constraint on the likely depth of Lake Arden. This is consistent with the decreasing spacing on ripples upwards through the lower sequence of silty mudstones. The medium grained, cross-bedded sandstones occur in thin sets of 0.2 to 0.5m inconsistent with a lake shoreline whilst small, parallel crested wave ripples are typical of very shallow water, in the order of a few centimeters. The jalodontid cranial remains were found in the lake shoreline deposits, having been transported and partly disarticulated in the lake shoreline swash zone.

### Systematic Paleontology

Class CHONDRICHTHYES Huxley, 1880

Subclass ELASMOBRANCHII Bonaparte, 1838

Order JALODONTIFORMES Ivanov, Duffin and Richter, 2021

Family JALODONTIDAE Ginter, Hairpetian and Klug, 2002

Genus KEUPERODUS Ivanov, Duffin and Richter, 2021

#### Type and only species

*Keuperodus brodiei* Ivanov, Duffin and Richter, 2021.

#### Revised diagnosis

Small teeth with a tricuspid crown; straight, short cusps ornamented with coarse, straight ridges diverging from the apex; central cusp is slightly shorter than lateral ones, with lanceolate ridges on labial face; thick tooth base directed lingually, elongated mesio-distally, strongly vascularized, bearing two shallow depressions on the occlusal side; numerous foramina on all faces of the base; the posterior of the basicranial surface bears two symmetrical transverse ridges that form a central groove pitted with several minute foramina. Basicranium with pronounced ventral angle. Basicranial circulation deeply embedded within the braincase with dorsal aorta bifurcating into the lateral dorsal aortae anterior to the level of the occiput. Internal carotids enter cranial cavity from within basicranium. Orbital arteries enter cranial cavity before passing into orbit. Palatoquadrate and Meckel’s cartilage elongate; palatoquadrate with low otic process.

*KEUPERODUS BRODIEI* (Ivanov, Duffin and Richter, 2021) (Figs. 3a, b – 7)

*Phoebodus brodiei* Woodward, 1893:281–287, pl. X, figs. 2–4 (original description).

**FIGURE 4.**
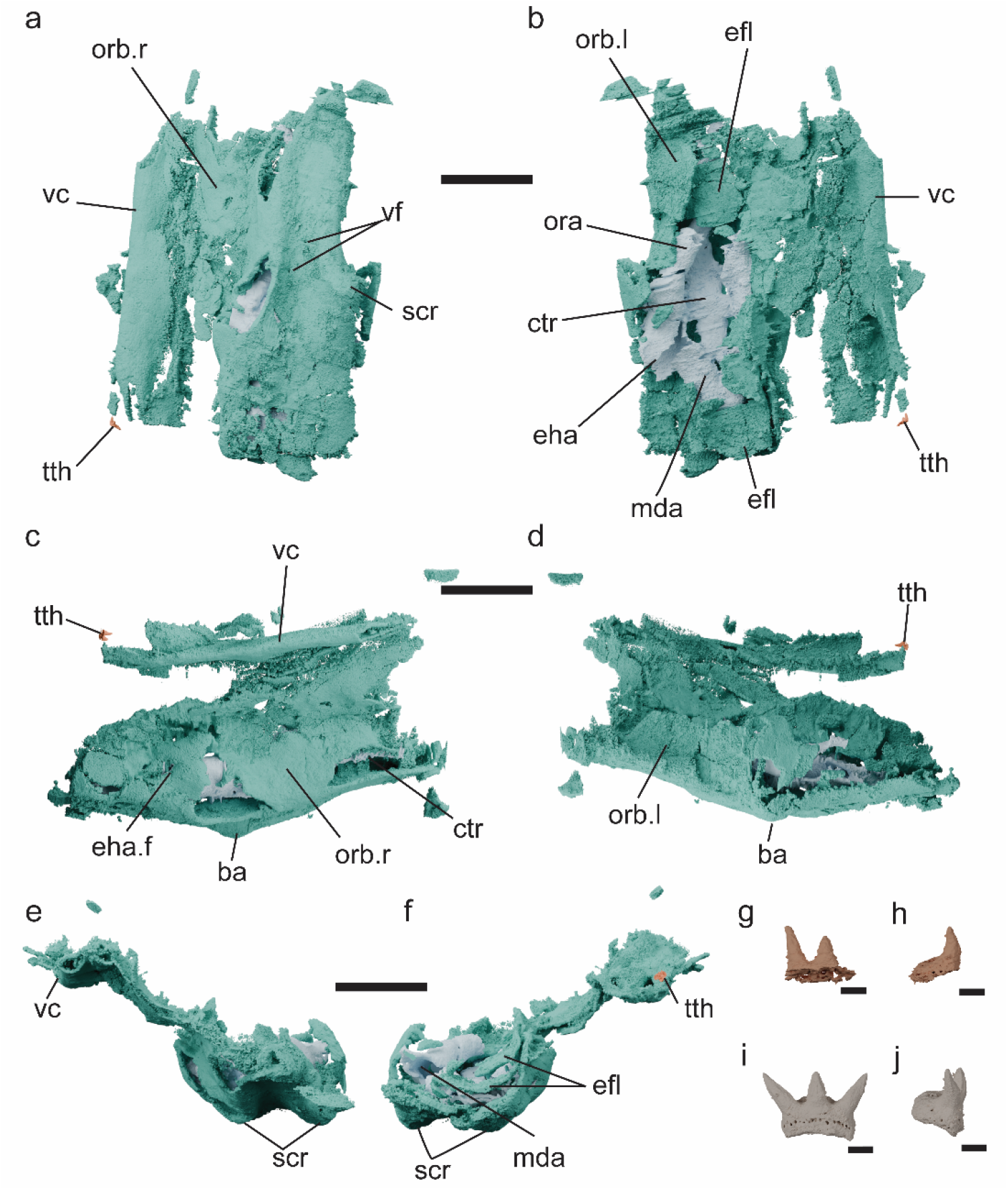
Partial braincase of *Keuperodus brodiei* visualized by CT imaging of WARMS G136: a, ventral view; b, dorsal view; c, dextral view; d, sinistral view; e, anterior view; f, posterior view; g,h, associated tooth in (g) labial view and (h) lateral view; i,j, tooth from *K. brodei* lectotype NHMUK PV P 7607 visualised by CT imaging in (i) labial view and (h) lateral view. Color scheme: turquoise, external neurocranium; light blue, basicranial canals; orange, tooth. Scale bar a-f 20 mm, g-j 1 mm.5. Abbreviations: ba, basal angle; ctr, canal for cerebral trunk; efl, endocranial floor; eha, canal for efferent hyoidean artery; eha.f, foramen for efferent hyoidean artery; mda, canal for median dorsal aorta; ora, canal for orbital artery; orb.l, orbital wall left; orb.r, orbital wall right; scr, subcranial ridges; tth, tooth; vc, visceral cartilage; vf, ventral foramina. (Full page width: 182.033 mm x 220 mm)

*Phoebodus brodiei* Woodward,1893: Cappetta, 1987:29, fig. 35; Cappetta, 2012:83–84, fig. 70A.

*Phoebodus keuperinus* Seilacher, 1948:24, figs. 1, 2.

*Phoebodus keuperinus* Seilacher, 1948: Cappetta, 2012:83, fig. 70B–G.

*Dittodus keuperinus* (Seilacher, 1948): Zidek, 1973:97.

*“Phoebodus” brodiei* Woodward, 1893: Ginter & Ivanov, 1996:264.

*“Phoebodus” keuperinus* Seilacher, 1948: Ginter & Ivanov, 1996:264.

*Keuperodus brodiei* Ivanov, Duffin & Richter, 2021:1 figs. 2-3.

#### Type Specimen

Lectotype, NHMUK PV P 7607, two associated tooth families, collected by Rev P.B. Brodie (Ivanov et al., 2021).

#### Type Locality and Stratigraphy

Cutting on the Grand Union Canal, Shrewley, Warwickshire, England. Arden Sandstone Formation, Upper Triassic, Carnian.

#### Diagnosis

Same as for genus.

#### Referred Material

In addition to the lectotype Ivanov et al. (2021) attributed eight isolated teeth to the species: NHMUK PV P 7608, NHMUK PV P 7609, NHMUK PV P 2266, NHMUK PV P 29483, NHMUK PV P 54639, NHMUK PV P 29484, SMNS 87883, and SMNS 88168-2. We also refer endoskeletal specimens WARMS G136 (Fig. 3a) and NHMUK PV P 7611 (Fig. 3b) to the species.

### Description

Ivanov et al. (2021) extensively described the two associated tooth families of *Keuperodus brodiei* collected from Shrewley that comprise the lectotype. We have refigured this specimen (Fig. 4i,j, Fig. S1) to facilitate comparison but do not redescribe it. Below we describe additional endoskeletal remains from Shrewley attributable to *K. brodiei*.

#### Keuperodus brodiei

WARMS G136 (Fig. 3a) preserves a part of the basicranial floor exposed in ventral view as well as a possible portion of a visceral cartilage and a single partial tooth (Fig. 4g-h). The partial tooth bears a tricuspid crown, though the third cusp is missing where it meets the edge of the matrix. Both preserved cusps are short, straight and diverge from the apex with the central cusp being slightly shorter than the lateral one (Fig 4g-h, Fig. S1k-p), and each cusp has a separate, elongate pulp cavity (Fig. S1q-s). The tooth base is thick, directed lingually, elongated mesio-distally and strongly vascularized (Fig. 4g-h). The resolution of the scan data is insufficient to establish whether ornament is present on the cusps. However the size and external and internal morphology of the tooth are consistent with the teeth preserved in the lectotype of *Keuperodus brodiei* NHMUK PV P 7607 (Fig. 4i-j, Fig. S1 b-j). Although the lateral cusps are less divergent than in most of the teeth in NHMUK PV P 7607, Ivanov et al. note that the teeth of the taxon are somewhat heterodont, and differences in cusp divergence are apparent in other jalodontid taxa such as *Jalodus australensis* (Ivanov et al. 2021, fig. 4)

The basicranial floor is slightly crushed dorsoventrally but still three-dimensional and broadly symmetrical. The basicranial surface is concave anteriorly with two distinct symmetrical ridges forming its ventrolateral edges. The gutter formed by this is punctured by two small foramina (Fig. 4a). Posteriorly these ridges flare laterally and ventrally to form a marked ventral angle. Posterior to this the basicranial floor is smooth and convex continuing until the edge of the rock with no evidence of having reached the occiput. Dorsally the basicranial floor continues to form the sides of the posterior orbit and anterior otic region: these have both slumped laterally (Fig. 4b). The postero-laterally oriented surface of the otic region is marked by a foramen. A straight, ridge of cartilage is preserved on the left-hand side of the specimen (Fig. 4a) that is interpreted here as a disarticulated fragment of the visceral cartilage compressed over the braincase rather than a portion of this braincase itself due to the absence of such linear cartilaginous features in other chondrichthyan braincases (Coates & Tietjen, 2017; Klug et al., 2023; Lane, 2010; Maisey, 2005; Maisey et al., 2019).

A complex and deeply invested series of canals for the basicranial circulation are preserved between the ventral surface of the braincase and a layer of cartilage that we interpret as the floor of the endocast (Figs 4b,e,f,5,6). These canals are somewhat crushed but largely intact and symmetrical with respect to the basicranial floor (Fig. 5). The dorsal aorta is enclosed by the basicranium at its most posterior extent and bifurcates into the left and right lateral dorsal aortae internally (Fig. 5b). Each of the lateral dorsal aortae has a short posterior divergence into the efferent hyoidean artery, which opened into the foramen in the otic wall (Fig, 4c). Ultimately, the passages for the lateral dorsal aorta extends anterodorsally to carry the orbital artery into the endocranial cavity. However, medio-ventrally they join twice to an unpaired median tube (possibly the hypophyseal chamber), that continues anteriorly. Two small foramina briefly open from this tube to the basicranial surface roughly halfway through its length. The artery continues to run toward the anterior extremity of the specimen, decreasing in width before bifurcating at the anterior edge of the specimen into the ophthalmic arteries (Fig. 5,6).

**FIGURE 5.**
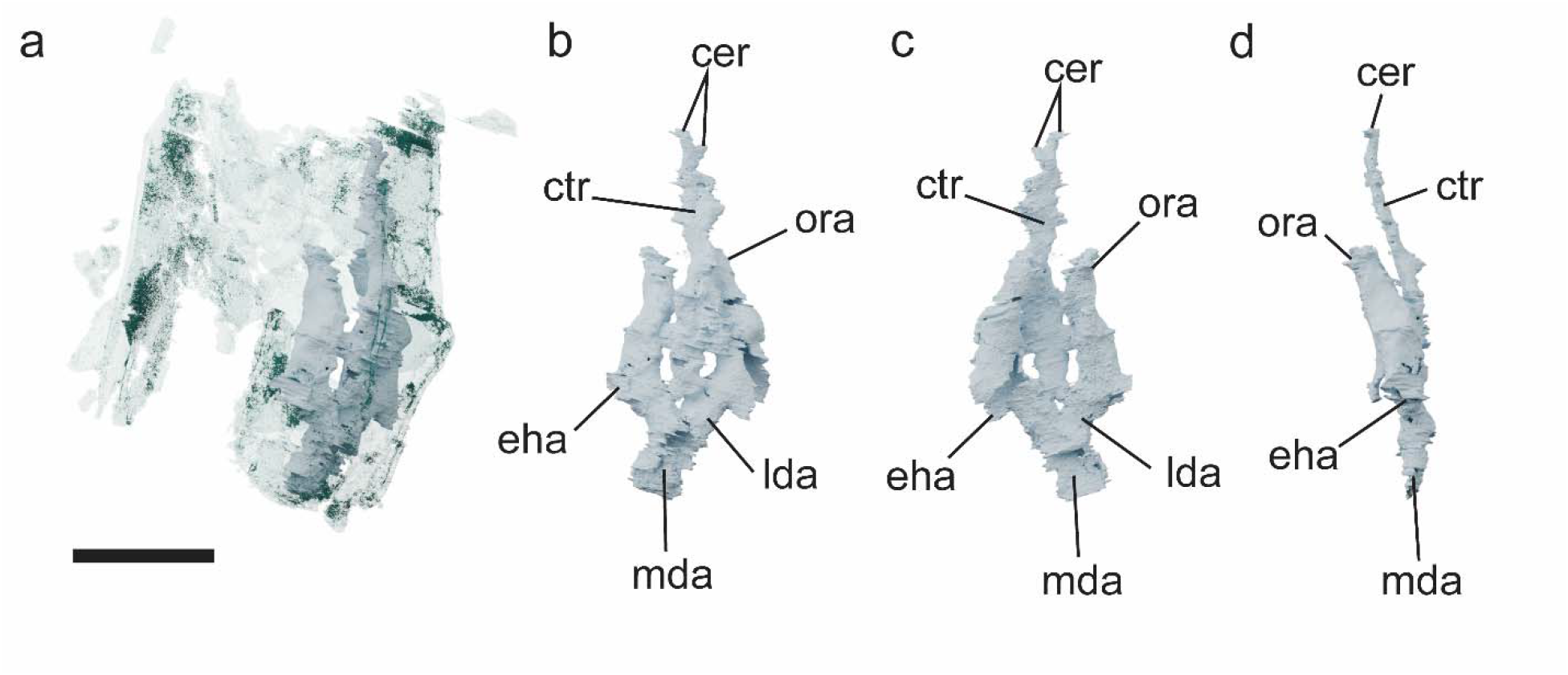
Basicranial circulation of *Keuperodus brodiei* visualized by CT imaging of WARMS G136: a, within the neurocranium, viewed from the matrix surface; b, ventral view; c, dorsal view; d, right lateral view. Scale bar 20 mm. Abbreviations: cer, cerebral carotids; ctr, canal for cerebral trunk; efl, endocranial floor; eha, canal for efferent hyoidean artery; eha.f, foramen for efferent hyoidean artery; ica, internal carotids; mda, canal for median dorsal aorta; ora, canal for orbital artery; scr, subcranial ridges. (Full page width: 182.033 mm x 78 mm)

**FIGURE 6.**
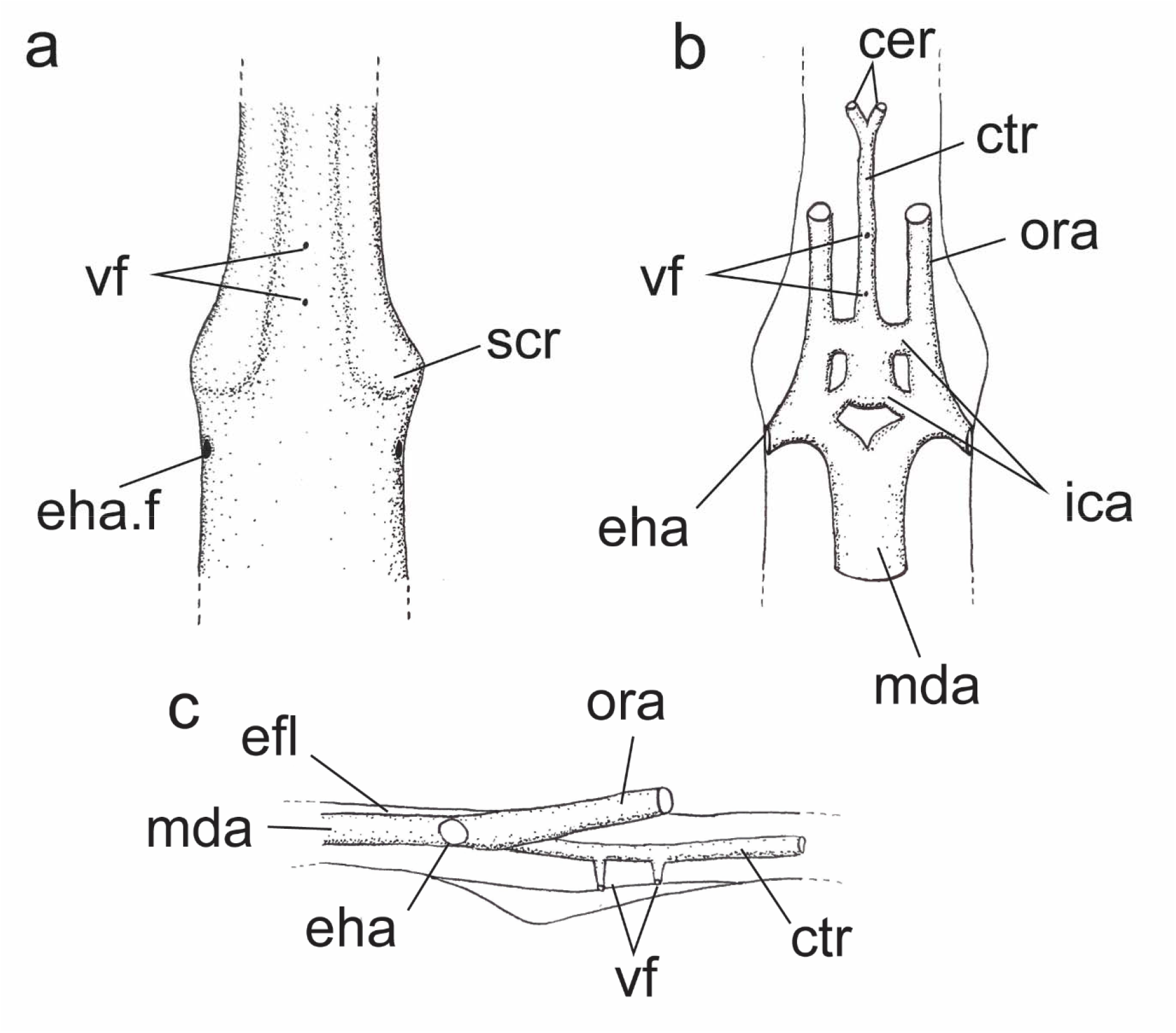
Reconstruction of braincase and basicranial circulation in *Keuperodus brodiei*: a, neurocranium in ventral view; b, basicranial circulatory network in ventral view; c, basicranial circulation network in lateral view. Abbreviations: cer, cerebral carotids; ctr, canal for cerebral trunk; eha, canal for efferent hyoidean artery; mda, canal for median dorsal aorta; ora, canal for orbital artery. (Single column width: 88.9 mm x 78 mm)

#### Keuperodus brodiei

NHMUK PV P 7611 preserves the posterior of the right palatoquadrate and Meckel’s cartilage in articulation, along with crushed pieces of cartilage that may be part of the hyoid arch and branchial rays, as well as a complex piece of cartilage that may be a crushed portion of the neurocranium but which we are unable to identify (Fig. 7 a-b). The palatoquadrate otic process is very low with a long diastema separating the articulation from the dental sulcus, and a pronounced extrapalatoquadrate ridge demarcating the dorsal border of the adductor fossa. The Meckel’s cartilage is well-rounded and elongated with a mandibular knob and quadrate condyle, and with a well defined postero-lateral ridge (Fig. 7 c-f).

**FIGURE 7.**
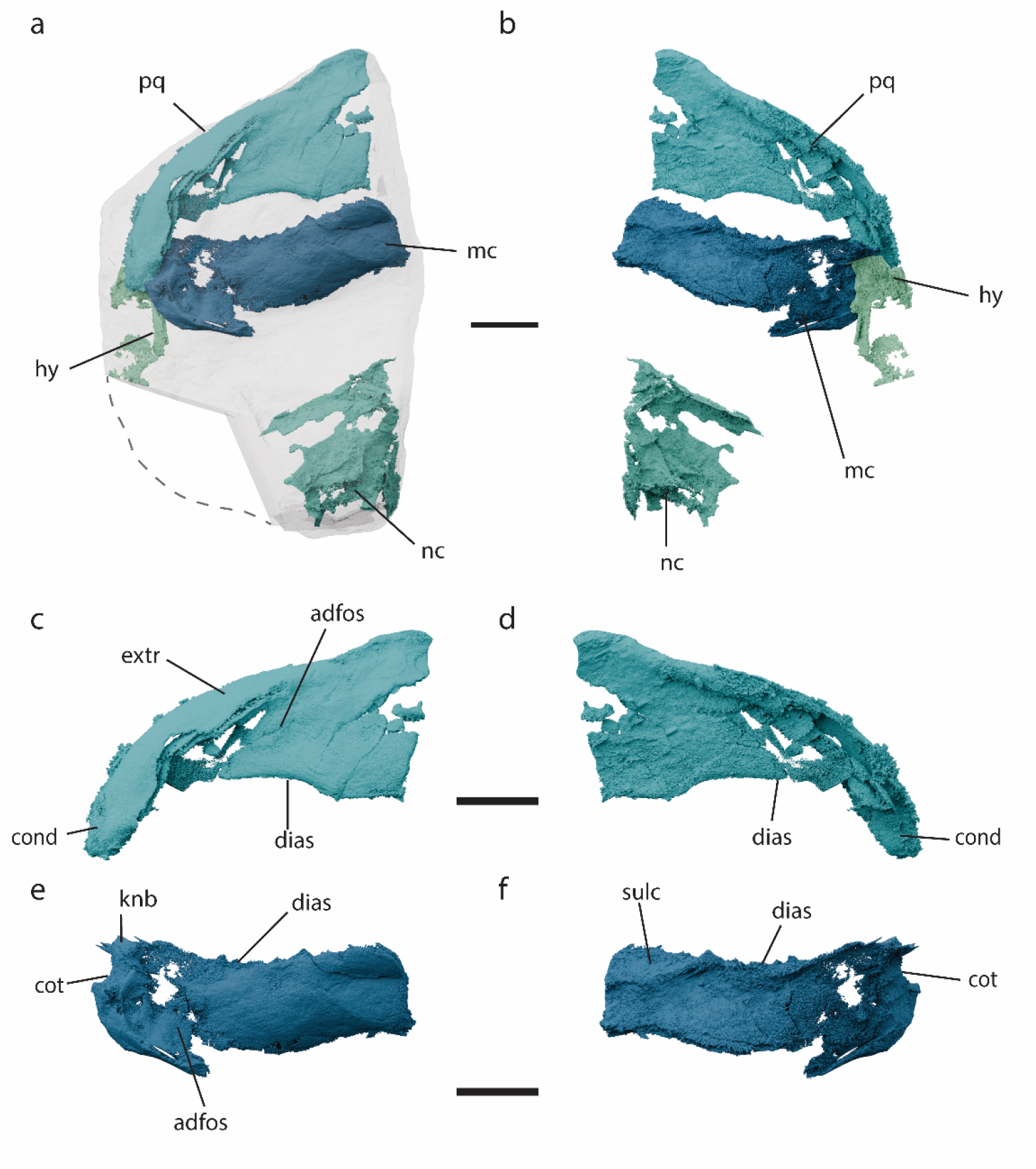
*Keuperodus brodiei* NHMUK PV P 7611, including parts of jaws and hyoid skeleton and a likely piece of braincase, visualized by CT imaging: a,b, scaled and stitched together models of jaws and braincase with jaws in right lateral view juxtaposed against matrix (a) and with jaws in left lingual view; c,d, right palatoquadrate in lateral (c) and lingual (d) view; e,f, right Meckelian cartilage in lateral (e) and lingual(f) view. Dashed line indicates approximate bounds of unscanned section. Scale bars 20 mm. Abbreviations: adfos, adductor fossa; cond, quadrate condyle; dias, diastema; extr, extrapalatoquadrate ridge; hy, hyoid fragments; knb, mandibular knob; mc, Meckelian cartilage; nc, neurocranium; pq, palatoquadrate; sulc, dental sulcus. (Full page width: 182.033 mm x 205 mm)

### Remarks

Only the teeth of *Keuperodus brodiei* and the hybodontid *Palaeobates keuperinus* have been recovered from Shrewley. Based on this and the similarity in its morphology to the lectotype (Ivanov et al. 2021) we assign the partial tooth in WARMS G 136 to *K. brodiei*.

Meanwhile the extensive embedding of the basicranial circulation in the neurocranium of WARMS G 136 is unlike that of any hybodont, and in particular *Acronemus tuberculatus* (Maisey 2011; Rieppel 1982), probably the closest comparison to *Palaeobates keuperinus* for which there is well-known neurocranial material. Based on this we assign WARMS G 136 to *Keuperodus brodiei* with confidence.

We assign NHMUK PV P7611 to *K. brodiei* based on its elongate jaw structure. This is not typical of Hybodontiformes and allied taxa (Fig. 8e,f) (Lane and Maisey 2012; Maisey, 1982; Romano and Brinkman, 2010), in particular *Acronemus tuberculatus* (Rieppel 1982) in which the jaws are short with a tall otic process (Fig. 8e), meaning that we consider it improbable that they belong to *Palaeobates keuperinus*. Instead they compare more closely to Phoebodontiformes (Fig. 8b,d)(Frey et al., 2019; Grogan & Lund, 2008; Klug et al., 2026). NHMUK PV P7611 was also collected from the same location by the same person, bears a nearly identical matrix to WARMS G136 and is of a similar relative size with the cartilage of WARMS G136, suggesting that it is possible they represent the same individual, although we are unable to establish this for certain. Brodie is known to have split specimens between what is now WARMS and the NHMUK, for example the holotype of the Bromsgrove Sandstone rhynchosaur *Langeronyx brodiei* (Ezcurra et al. 2016).

**FIGURE 8.**
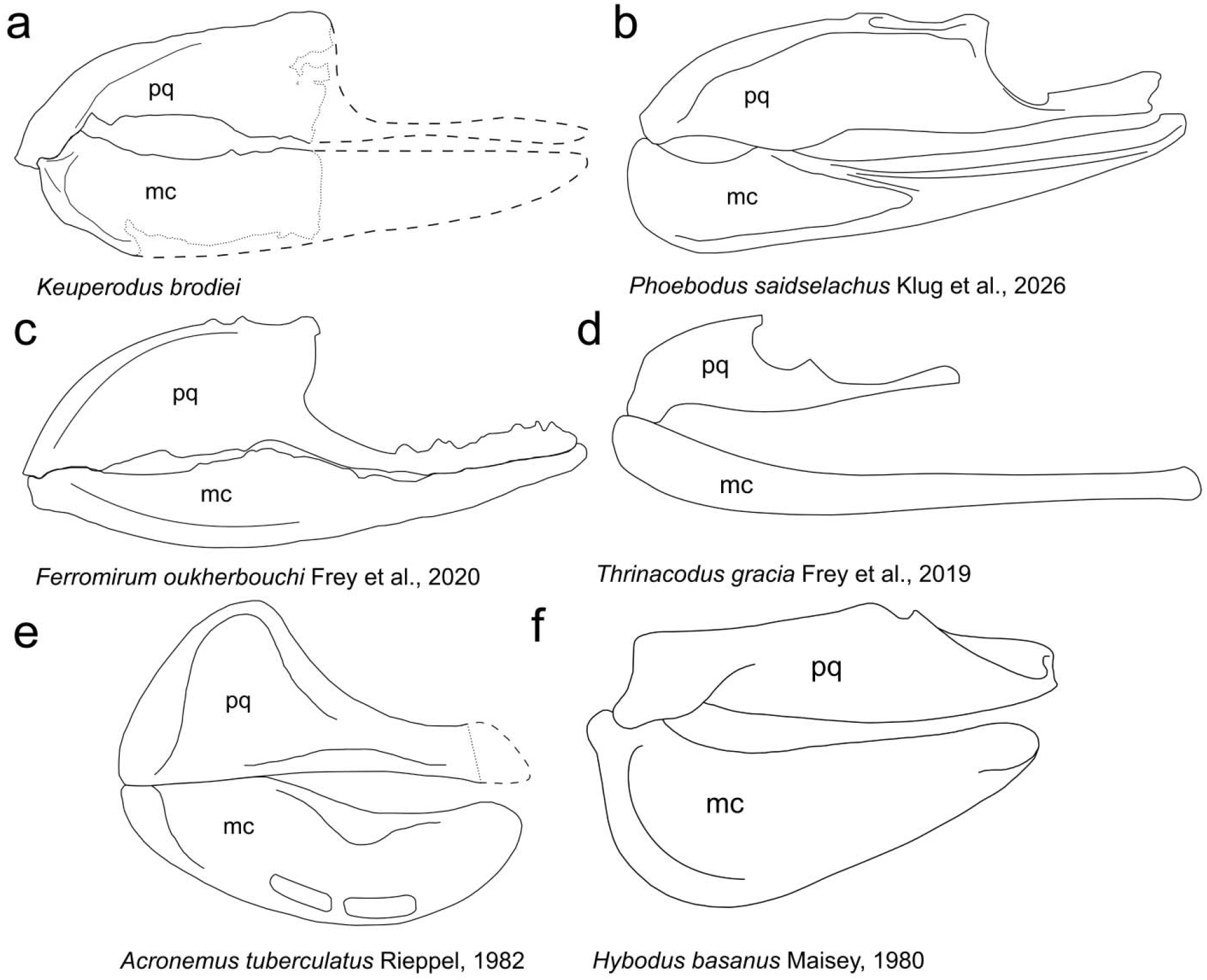
Drawings of the jaw cartilages of various chondrichthyan taxa compared to Keuperodus brodiei: a, Keuperodus brodiei; b, Phoebodus saidselachus; c, Ferromirum oukherbouchi; d, Thrinacodus gracia; e, Acronemus tuberculatus; f, Hybodus basanus. (Two-thirds page width: 122 mm x 100 mm)

Subclass ELASMOBRANCHII Bonaparte, 1838 *incertae sedis*

#### Description

NHMUK PV P 7612a-d (Fig. 3b-f) comprises pieces of articulated postcranial skeleton. NHMUK PV P 7612a and NHMUK PV P 7612b both preserve a portion of the dorsal fin basal cartilage; both specimens also contain one basal fragment of a fin spine in articulation (Figs 3c,d, 9a-d). The specimens do not articulate with each other; hence, one is possibly representative of the anterior dorsal fin and the other the posterior dorsal fin, though it is not possible to tell which is which. NHMUK PV P 7612b (Figs 3d, 9c,d) preserves a small section of striated ornament on the dorsal tip of its fin spine fragment (Fig. 3c) NHMUK PV P 7612c preserves an articulated row of eight flat elements with a marked kink one quarter of the way along their length (Figs 3e, 9e-f), which we consider likely to be caudal fin hypurals based on comparison with other chondrichthyans (Coates & Sequiera, 2001; Maisey, 1983). NHMUK PV P 7612d preserves what are probably pectoral fin radials (Figs 3f, 9g-h).

**FIGURE 9.**
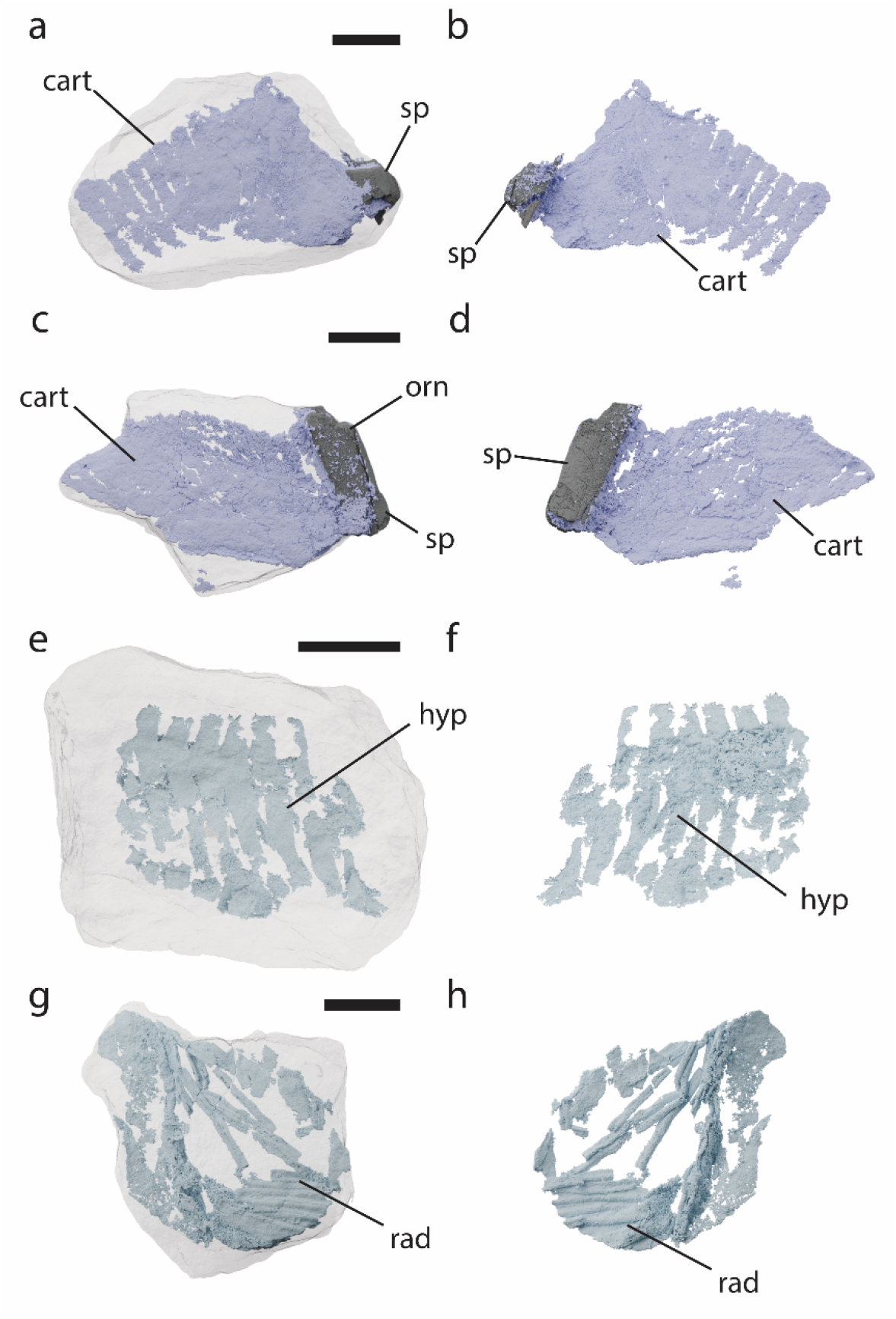
Postcranial remains Elasmobranchii *incertae sedis* from Shrewley NHMUK PV P 7612a-d visualized by CT imaging: a, b, NHMUK PV P 7612a dorsal fin spine and cartilage in right lateral view juxtaposed against a semi-transparent matrix (a), and left lateral view (b); c,d, NHMUK PV P 7612b dorsal fin spine and cartilage in right lateral view juxtaposed against semi-transparent matrix (c), and left lateral view (d); e,f, NHMUK PV P 7612c hypurals in left lateral view juxtaposed against semi-transparent matrix (e), and right lateral view (f); g,h, NHMUK PV P 7612d fin radials in left lateral view juxtaposed against semi-transparent matrix (g), and right lateral view (h). Scale bars 20 mm. Abbreviations: cart, dorsal fin spine cartilage; hyp, hypurals; orn, ornament patch; sp, fin spine; rad, radials. (Two-thirds page width: 122 mm x 180 mm)

#### Remarks

We place these specimens in Elasmobranchii *incertae sedis* due to the lack of phylogenetically diagnostic characters. It is possible that these pieces may belong to one individual, and could have been collected from the same specimen as WARMS G136 and NHMUK PV P 7611 due to the close similarity in host matrix and the collection history. However, the striated ornament resembles complete isolated fin spines from the Arden Sandstone Formation assigned to *Palaeobates keuperinus* (Murchison & Strickland, 1837; Woodward, 1899a). It is unclear whether this ornament is diagnostic or whether the attribution of these isolated fin spines exclusively to *Palaeobates* is accurate.

### Proposed interpretation of basicranial circulation in *Phoebodus saidselachus*

The neurocranium of *K. brodiei* bears similarities with that of the only three-dimensionally preserved phoebodontid neurocranium, the Upper Devonian *Phoebodus saidselachus* (Frey et al., 2019; Klug et al., 2026) (Fig. S2). In *P. saidselachus* two canals parallel to the notochordal canal are identified as having carried the glossopharyngeal (IX) nerve by Klug et al. (2026, fig. 9). Although interpreting these is hindered by a break in the specimen, the preserved anatomy of these paired canals is inconsistent with carriage of the glossopharyngeal (IX) nerve: they do not connect with the intracranial space via the labyrinth, are ventrally and medially displaced from the labyrinth (Fig. S2c,d,f), have smaller ramules branching dorsally (Fig. S2c,d), and do not exit the neurocranium through the posterior opening identified by Klug et al. (2026) as the glossopharyngeal (IX) foramen (Fig. S2a). Instead, we consider it more likely that a posterior extension of the base of the saccular chamber that aligns with the glossopharyngeal foramen carried the glossopharyngeal (IX) nerve out of the neurocranium (Fig. S2a,b,f), similar to the arrangement in Paleozoic elasmobranchs such as *Tamiobatis vetustus* (Maisey, 2026).

We consider it more plausible that the canals identified by Klug et al. (2026) as having carried the glossopharyngeal (IX) nerve are instead the posterior sections of deeply embedded lateral dorsal aortae. This is supported by the alignment of the canals with anterior parts of deeply embedded neurocranial vasculature, their probable posterior alignment with the paired foramina on either side of the occiput identified as entry points for the lateral dorsal aorta in *P. saidselachus* by Frey et al. (2019, fig. 2), and the dorsal ramules, which are likely to be musculospinal arteries passing into the glossopharyngeal-vagal passage as is common in Paleozoic elasmobranchs (Maisey, 2026. Fig. 10). Anteriorly, these lateral dorsal aortae are aligned with a series of canals that are not interpreted in detail by Klug et al. (2026) but which we interpret as anterior parts of the basicranial circulation having similarities with the layout of these blood vessels in other vertebrates (Rahmat & Gilland, 2014). The lateral dorsal aortae join medially into a united tube either for the internal carotids or an unpaired carotid trunk, which then continues anteriorly before entering a sub-hypophyseal space that is open ventrally through a basal fenestra. Here it splits into two cerebral carotids, giving rise to the ophthalmic arteries and entering the cranial cavity anterior to the hypothalamus (Fig. S2). Notably, although on the limits of the data’s resolution this single internal carotid canal appears to have two small rami passing ventrally through the basicranial floor. The consequences of this reinterpretation are that *P. saidselachus* had a basicranial circulation that was unusually extensively embedded in the basicranium and which, like that of *K. brodiei* included an unpaired internal carotid canal that passed ventral to the hypothalamus.

**FIGURE 10.**
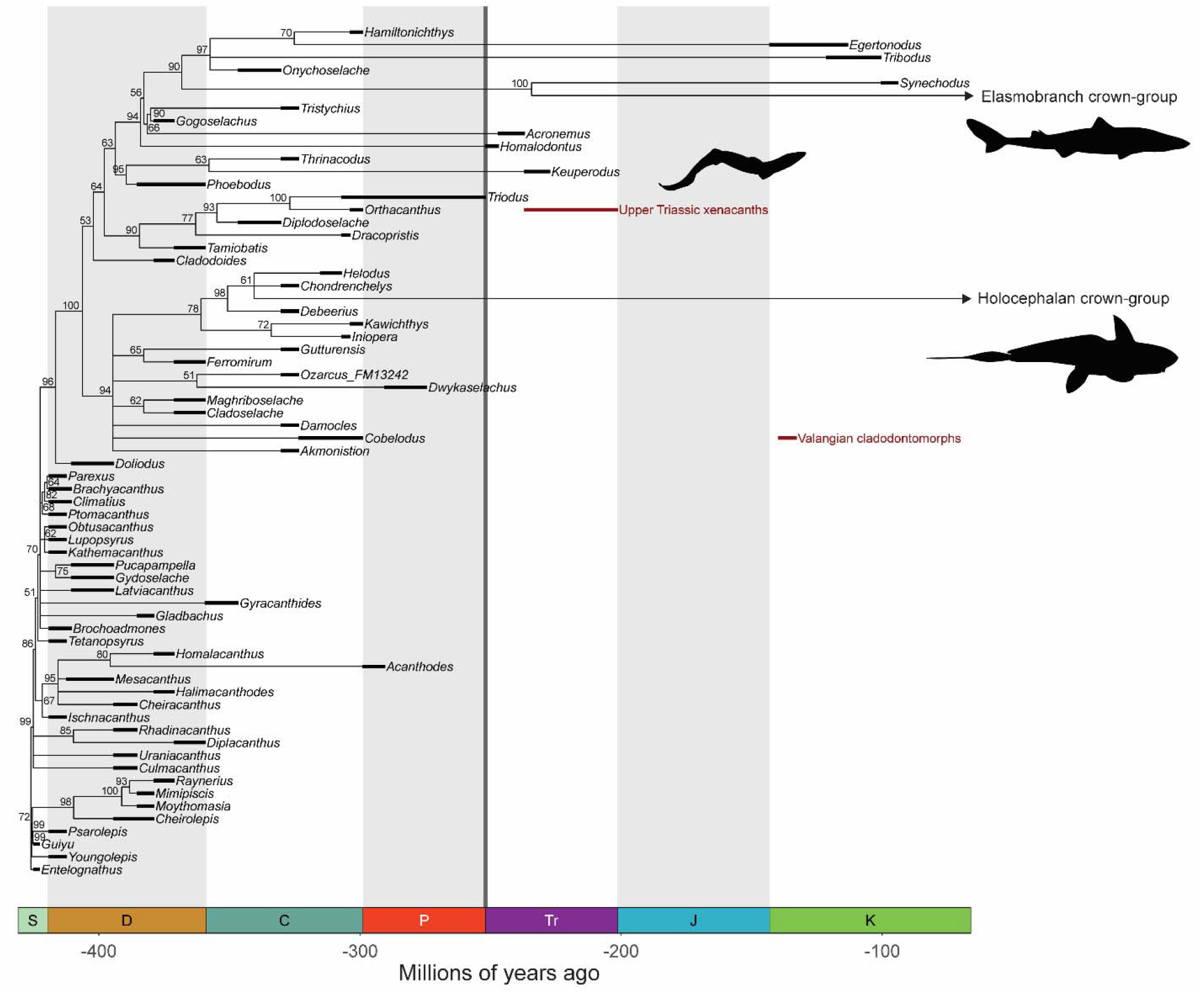
Time-scaled results of phylogenetic analysis plotted against time, including late dental records for “Paleozoic” chondrichthyans. Tree shown is the 50% majority rule consensus tree resulting from the Bayesian analysis, with numbers representing posterior probabilities of nodes. Grey line represents Hangenberg extinction. Tree shown is the scaled *a posteriori* using tip ages (see supplementary table 2) and the timePaleoPhy function in paleotree package for R. Abbreviations: C, Carbonferous; D, Devonian; J, Jurassic; K, Cretaceous; P, Permian; S, Silurian; Tr, Triassic. (Full page width: 182.033 mm x 150 mm)

### Phylogenetic Analysis

The parsimony analysis recovered 96 most-parsimonious trees (MPTs) with 582 steps. All trees had a CI of 0.428 and an RI of 0.756, indicating moderate to low homoplasy in the dataset. In both the parsimony strict consensus of these trees and the Bayesian majority rule consensus of a posterior sample of 7500 trees *Keuperodus brodiei* is recovered as a stem-group elasmobranch, in a clade with *Thrinacoselache* and *Phoebodus* (Figs 10, S3). In the parsimony strict consensus the three taxa are recovered in a polytomy, in the Bayesian tree the clade including the three taxa has a posterior probability of 95% and *K. brodiei* and *Thrinacodus* are recovered as sister-taxa to the exclusion of *Phoebodus* with a posterior probability of 63%. In the parsimony strict consensus this clade is recovered in a polytomy with other members of the elasmobranch total group, in the Bayesian consensus it is the sister group to crown-chondrichthyans and hybodonts to the exclusion of xenacanths, ctenacanths and *Cladodoides*. The broader topology of these trees otherwise reflects that recovered by Klug et al. (2026).

## Discussion

### Phylogenetic relationships of Jalodontiformes

Before Ivanov et al.’s (2021) establishment of the Order Jalodontiformes, *Keuperodus brodiei* teeth from Shrewley were attributed to a Family within the Order Phoebodontiformes due to their tricuspidate morphology (Ginter et al., 2010). However, Ivanov et al. (2021) argued a closer similarity to jalodontids such as *Adamantina* based on characters including relatively robust cusps without sigmoidal flexion and pronounced ornamentation and erected a distinct Order for jalodontid teeth. Here we provide the first information on the cranial morphology of Jalodontiformes, which suggests skeletal similarities between *K. brodiei* and phoebodontids. Like *Phoebodus saidselachus* (according to our interpretation above), much of the basicranial circulation including the lateral dorsal aortae are embedded in the basicranium and both the palatoquadrate and Meckel’s cartilage have an elongated form that is seen in *Phoebodus* and *Thrinacoselache* (Fig. 8b,d) (Frey et al., 2019; Grogan & Lund 2008; Klug et al., 2026) and to a certain extent in Symorriiformes like *Ferromirum* (Fig. 8c), in contrast to the shorter more robust jaws of hybodontiforms like *Hybodus* or allied Triassic taxa like *Acronemus* (Fig. 8e,d).

Our phylogenetic analyses finds evidence for a relationship between Jalodontiformes and Phoebodontiformes based on the tricuspidate tooth (character 66) and the deep embedding of the basicranial circulation. This provides evidence that jalodontids could have been derived from phoebodontids early in their evolution. Although the teeth of these taxa are morphologically distinct (Ivanov et al., 2021), similarities in cranial cartilages suggest a possible speciation event between the two during the Devonian. This is supported by an increase in origination rates in chondrichthyans during the Frasnian of the Upper Devonian (Schnetz et al., 2024). Ultimately, despite these new and unexpected similarities, the lack of fossil material known at present makes this hypothesis impossible to prove, and, as such, the family Jalodontidae should remain within Jalodontiformes until further jalodontid material is discovered that can confidently prove otherwise.

### Comparative basicranial anatomy

The arrangement of the basicranial circulation in *Keuperodus brodiei* combines features of the basicranial circulation from different chondrichthyan groups. In extant elasmobranchs the basicranial circulation forms a characteristic bell-shaped circuit external to the braincase with the dorsal aorta splitting into lateral dorsal aortae posterior to or level with the occiput and entering the braincase through one or more internal carotid foramina (Lane, 2010; Maisey, 1983), while in extant holocephalans it splits anterior to the occiput and does not enter the braincase medioventrally(Pradel et al., 2021). In hybodonts such as *Egertonodus basanus* and *Tribodus limae* (Lane 2010; Maisey, 1983), as well as in allied taxa such as *Acronemus tuberculatus* and *Tristychius arcuatus* (Coates & Tietjen, 2017; Maisey, 2011) the lateral dorsal aortae split posterior to the occiput. This is a common condition more generally in a broad range of Paleozoic chondrichthyans, being present in *Cladodoides*, *Tamiobatis, Doliodus*, *Pucapampella*, *Carcharopsis*, *Orthacanthus*, (Bronson et al., 2018; Maisey, 2005; Maisey, 2026; Maisey et al., 2009, 2019; Schaeffer, 1981) as well as in *Phoebodus* (Klug et al., 2026). Contrastingly in symmoriiforms such as *Dwykaselachus oosthuizeni* (Coates et al., 2017) and *Akmonistion zangerli* (Coates & Sequiera, 2001), the dorsal aorta also divides into lateral dorsal aortae anterior to the occiput level. In this sense *K. brodiei* resembles symmoriiforms more closely than other Paleozoic or Mesozoic chondrichthyans, although the dorsal aortae bifurcate at a markedly anterior level even compared to Symmoriiformes.

Another sense in which *K. brodiei* is unusual is that the basicranial circulation is embedded deeply and entirely within the basicranium with the lateral dorsal aortae never exiting onto the basicranial surface (Fig. 6). This contrasts with the condition in extant elasmobranchs, holocephalans and *Synechodus dubrisiensis*, in which none of the basicranial circuit is housed in the basicranium (Maisey, 1985; Pradel et al., 2021).

In *Egertonodus basanus*, *Cladodoides*, *Tamiobatis*, *Orthacanthus* only stretches of the lateral dorsal aortae are invested in the basicranium although in *Tribodus limae* a substantial proportion of the basicranial circuit is invested (Lane, 2010; Maisey, 2005; Schaeffer, 1981). In symmoriids, such as *Cobelodus*, *Ozarcus*, *Akmonistion* and *Dwykaselachus*, the median dorsal aorta and its split into lateral dorsal aortae are typically invested in the basicranium as are other sections of the circuit, but some stretches remain exposed (Coates et al., 2017; Maisey, 2007). The closest comparisons to *Keuperodus* are *Phoebodus* (based on our reinterpretation) and *Carcharopsis* in which the lateral dorsal aortae are extensively embedded in the basicranium, although in both taxa the lateral dorsal aortae split before entering the neurocranium (Bronson et al., 2018; Klug et al., 2026). In *Phoebodus* the entire basicranial circulation is invested in the basicranium (Klug et al., 2026). Similarly, in the Carboniferous *Carcharopsis* the lateral dorsal aortae are invested in the neurocranium with the orbital arteries extending directly from them into the orbital floor (Bronson et al., 2018). This extensive embedding of the basicranial circulation may represent a character state mostly present in Paleozoic taxa which was lost in lineages more closely related to the crown-group (Maisey, 2026), but which is obscured by poor sampling of endoskeletal remains.

### Paleozoic shark survivors of the EPME

Chondrichthyan phylogeny plotted against time illustrates that several distantly related groups survived the EPME (Fig. 10). Crown-group chondrichthyans may have diverged in the Permian (Marion et al., 2024) and so it is possible that the crown-group itself includes multiple successful traversals of the EPME. At least one lineage representing hybodontiforms and closely related taxa such as *Acronemus* also survived the extinction. However, the results of our phylogenetic analysis place *Keuperodus brodiei* in a clade relatively remote from the elasmobranch crown-lineage and these closely related EPME survivors (Fig. 10). Several groups of chondrichthyans characteristic of Paleozoic faunas survived the EPME into the Triassic alongside lineages closely related to modern crown-groups (Koot, 2013). Xenacanthiformes, a typically freshwater group of probable stem-elasmobranchs (Coates et al., 2017), survived into the Upper Triassic (Bhat et al., 2018; Hodnett et al., 2022) and are found in British Carnian deposits similar to those at Shrewley (Dawnson et al., 2022; Johnson, 1980). The enigmatic scale-taxon *Listracanthus* is found in a marine setting in the Lower Triassic, as are likely stem-holocephalan eugeneodontids (Mutter & Neuman, 2006, 2008; Mutter & Rieber, 2005). Cladodontomorph and ctenacanthiform shark teeth have been described from a deepwater marine setting in the Lower Cretaceous (Guinot et al., 2013). This places *Keuperodus brodiei* in context as one among a phylogenetically diverse set of Paleozoic chondrichthyan survivors of the EPME.

*Keuperodus brodiei* from the Arden Sandstone Formation of the English Midlands is a further example of a Triassic chondrichthyan with demonstrable Paleozoic characteristics found in a freshwater setting. The Carnian Grabfeld Formation of central and southern Germany, the only other locality where *K. brodiei* has been found (Ivanov et al., 2021) is interpreted as representing saline lakes surrounded by evaporitic mudflats along a coastal playa (Leitner et al. 2023). This makes *K. brodiei* unique amongst Jalodontiformes; all other jalodontid taxa are known from marine deposits (Ivanov et al., 2021). The Arden Sandstone Formation lacustrine depositional system formed during the intense rainfall associated with the Carnian Pluvial Episode and the establishment of a large evaporitic freshwater lake across the English Midlands that persisted throughout the Carnian Stage (Burley et al., 2023; Marshall, 2019; Simms & Ruffell, 2018). There is, however, evidence that freshwater ecosystems were affected equally as hard and potentially worse than marine ecosystems during the EPME (Mays et al., 2021). The presence of shark fin spines in the freshwater, Middle Triassic Otter Sandstone Formation of Devon, does provide direct evidence that post EPME, by 245 Ma, chondrichthyans were adapted to freshwater ecosystems in the UK (Coram et al., 2019). During the EPME, lacustrine and fluvial environments may have acted as refugia during times of marine crises (Wen et al., 2022), analogous to the role proposed for deepwater habitats and cladodont chondrichthyans (Guinot et al., 2013).

## Conclusions

Our sedimentological interpretation of the Arden Sandstone Formation shows that *Keuperodus brodiei* lived in the large freshwater Lake Arden, which had deepened and expanded after the onset of the Carnian Pluvial Episode. *Keuperodus brodiei* provides a clear example of long-term Paleozoic chondrichthyan survivorship across the EPME, with well-established freshwater ecosystems possibly acting as refugia. The rarity of such fossil material illustrates that the Arden Sandstone Formation at Shrewley provides a unique source and site of Triassic chondrichthyan skeletal material of international significance and along with recent work from Velin et al. (2026), suggests further exploration of Upper Triassic freshwater deposits could yield further calcified cartilage fossil material. The discovery of these specimens reinforces the need for collaboration between researchers and museums to re-examine long-established paleontological collections for unrecognized, scientifically important fossil material. Meanwhile the unusual morphology of the neurocranium revealed by this first description of endoskeletal material attributable to *K. brodiei* highlights the scarcity of sampling of chondrichthyan skeletal remains relative to the tooth fossil record particularly in the Triassic, and demonstrates the importance of these fossils to place tooth-based chondrichthyan taxa into a phylogenetic context.

## Acknowledgements

We thank Emma Bernard (NHMUK) for allowing the authors access to the NHMUK’s fossil fish collections. Many thanks to Agnese Lanzetti and Brett Clark (NHMUK) for scanning the NHMUK specimens and Sarah Jamison-Todd and Luke Meade (University of Birmingham) for scanning WARMS G136 as well as the NHMUK and the University of Birmingham for access to their CT scanners. We would also like to thank the Canal and River Trust for allowing access to the Shrewley site on the Grand Union Canal, and the Warwickshire Geological Conservation Group, whose geoconservation efforts greatly help preserve the few remaining outcrops of the Arden Sandstone Formation.

We are also grateful to Mike Coates, John Maisey, Alan Pradel, and Christian Klug for valuable conversations regarding the interpretation of the fossil specimens in this study. Thanks to Sam Giles for access to segmentation software and hardware. RPD is funded by the National Environment Research Council Independent Research Fellowship [grant number UKRI4180].

## Supplementary tables

**Supplementary table 1:** Details of CT scanning for specimens in this study.

| **Fossil Specimen** | **Scan Specifications** |  |  |  |  |  |
|---|---|---|---|---|---|---|
|  | kV | µA | Voxel size (µm) | Cu Filter (mm) | Exposure (ms) | No. of Projections |
| WARMS G136 | 190 | 165 | 31.95 | 1.5 | 1000 | 4492 |
| NHMUK PV P 7611 (targeting jaws) | 197 | 149 | 60.36 | 2 | 354 | 3500 |
| NHMUK PV P 7611 (targeting neurocranium) | 197 | 155 | 34.26 | 2 | 500 | 0 |
| NHMUK PV P 7612a | 170 | 149 | 51.14 | 1 | 354 | 3500 |
| NHMUK PV P 7612b | 170 | 149 | 44.61 | 1 | 354 | 3500 |
| NHMUK PV P 7612c | 170 | 149 | 41.49 | 1 | 354 | 3500 |
| NHMUK PV P 7612d | 170 | 149 | 36.51 | 1 | 354 | 3500 |

## Supplementary Figures

**SUPPLEMENTARY FIGURE 1.**
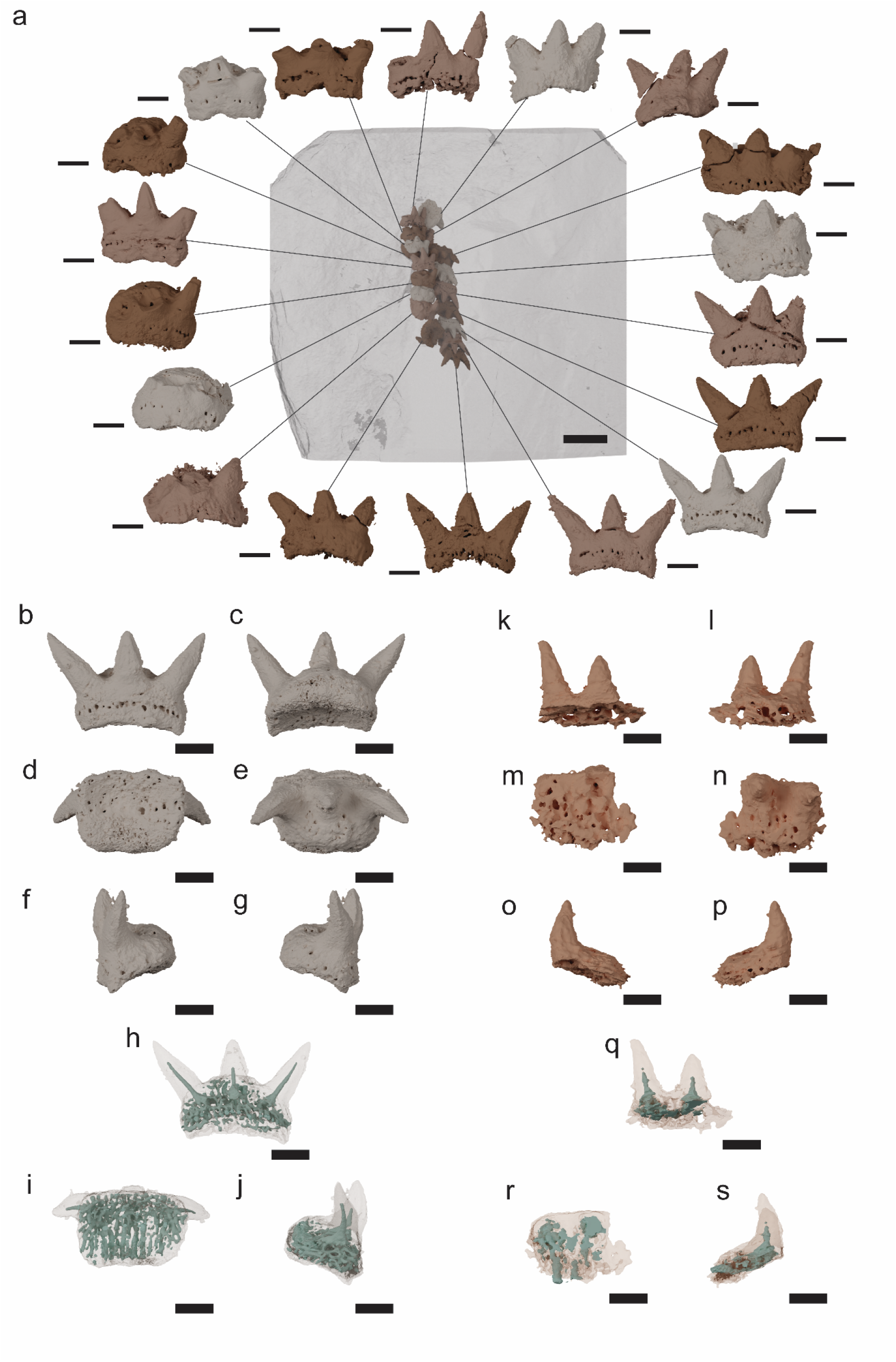
*Keuperodus brodiei* lectotype NHMUK PV P 7607 visualised using CT dataset from Ivanov *et al*. 2021: a, entire specimen with rock matrix rendered transparent and individual teeth shown in labial view; b-g, tooth from bottom right corner of panel a shown in (b) labial view, (c) lingual view, (d) ventral view, (e) dorsal view, (f) sinistral view, (g) dextral view; h-j, same tooth shown with tissue rendered semi-transparent and internal vasculature in blue in (h) labial view, (i) ventral view, and (j) dextral view; k-p, tooth from WARMS G 136 shown in (k) labial view, (l) lingual view, (m) ventral view, (n) dorsal view, (o) sinistral view, (p) dextral view; q-s, same tooth shown with tissue rendered semi-transparent and internal vasculature in blue in (q) labial view, (r) ventral view, and (s) dextral view. Scale bars 1 mm except for scale bar overlying matrix in panel a which is 5 mm.

**SUPPLEMENTARY FIGURE 2.**
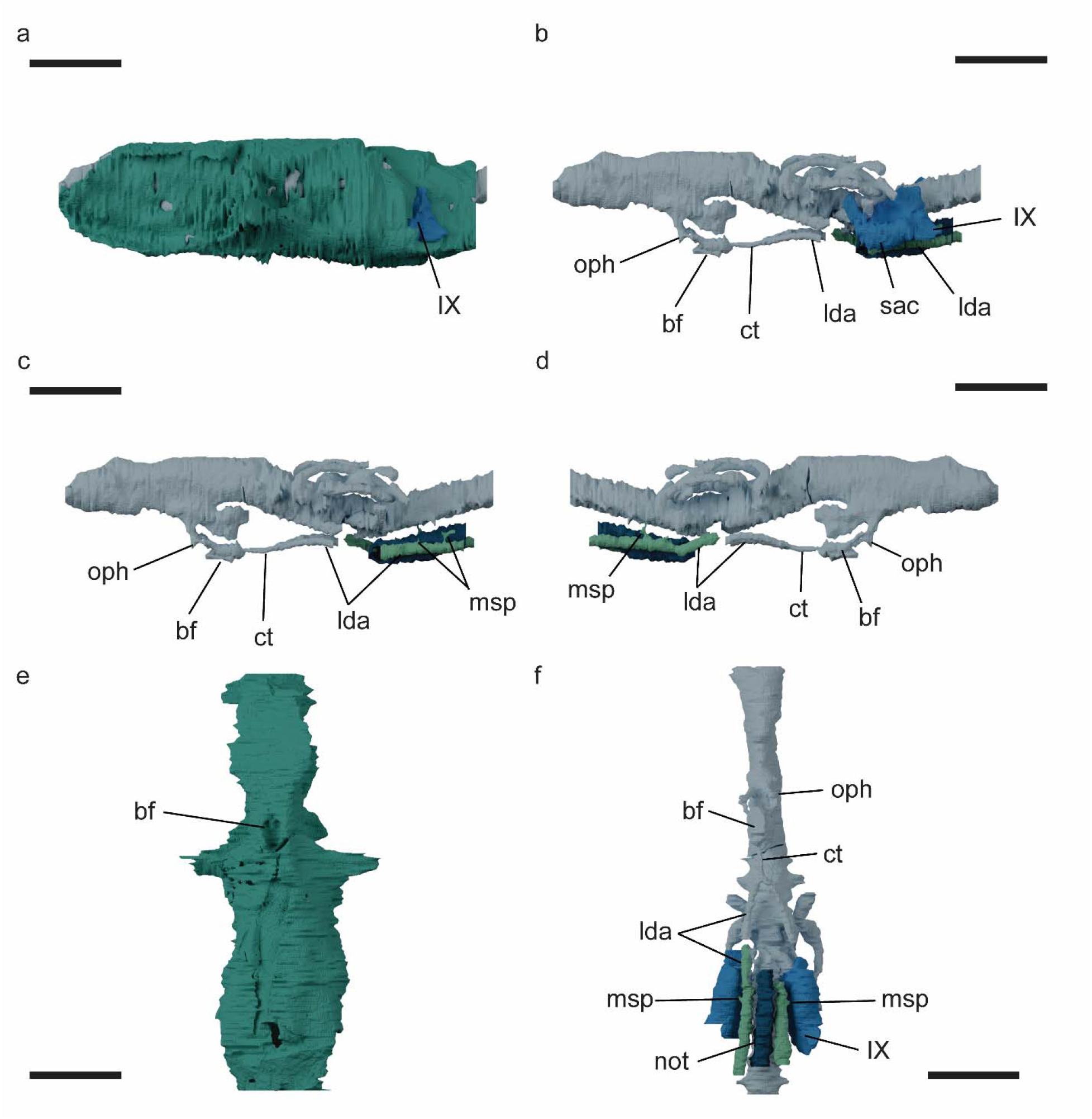
Reinterpretation of the basicranial circulation of *Phoebodus saidselachus* using renders of models from Klug et al. (2026): a, neurocranium in left lateral view with endocast; b, endocast in left lateral view; c, endocast in left lateral view with saccular chamber and glossopharyngeal canal removed; d, endocast in right lateral view with saccular chamber and glossopharyngeal canal removed; e, neurocranium in ventral view; f, endocast in ventral view. Colors: turquoise, neurocranium; light blue, endocranium; blue, saccular chamber and glossopharyngeal canal; green, lateral aortica canals; dark blue, notochordal canal. Scale bars 50 mm. Abbreviations: bf, basicranial fenestra; ct, carotid trunk; lda, lateral dorsal aorta; msp, dorsal ramules carrying musculospinal arteries; not, notochordal space; oph, ophthalmic artery; sac, saccular chamber; IX, glossopharyngeal foramen/inferred route of nerve.

**SUPPLEMENTARY FIGURE 3.**
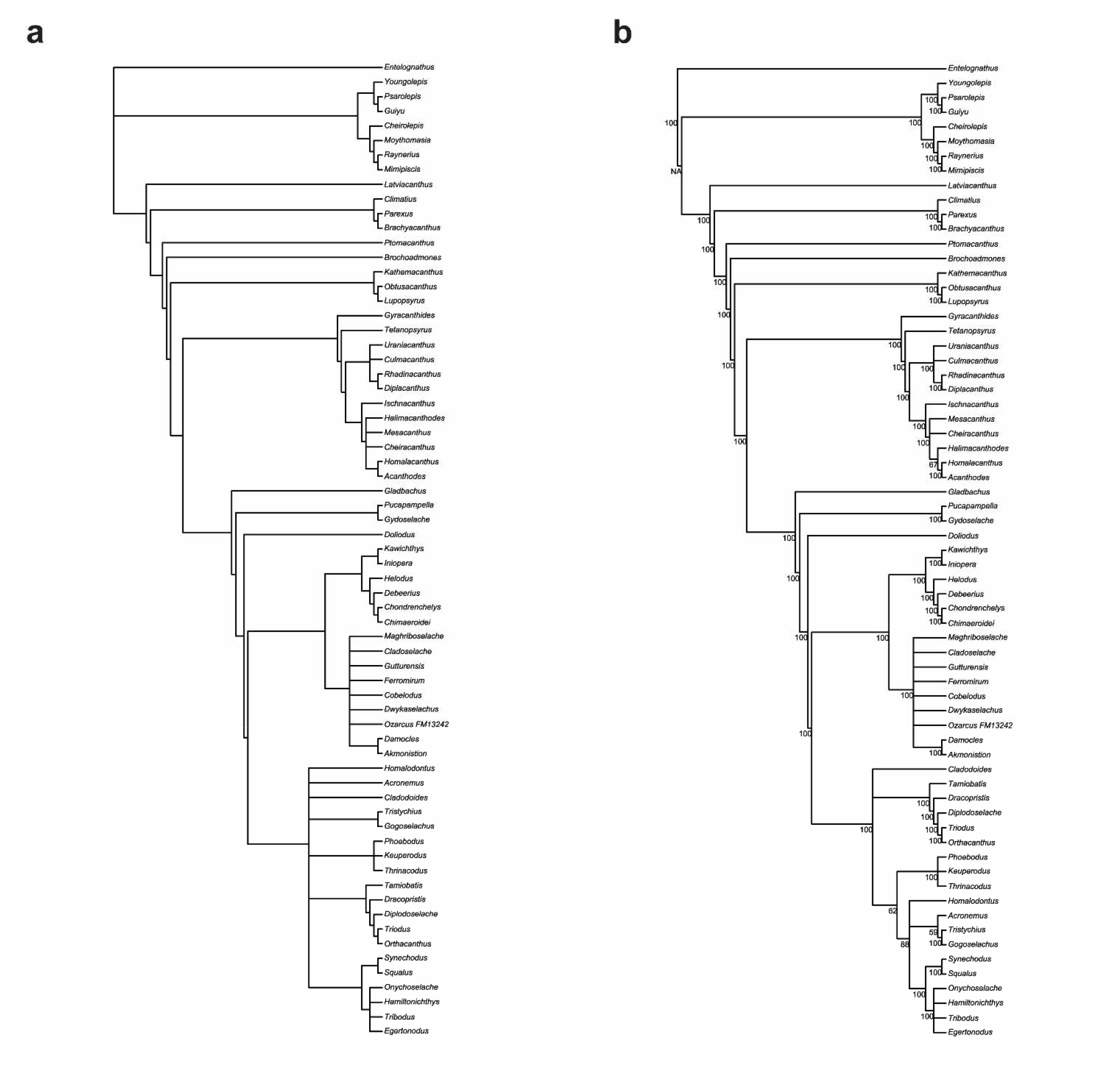
Results of the parsimony analysis: a, strict consensus tree; b, 50% majority rule consensus tree.

## Notes

### Competing Interest Statement

The authors have declared no competing interest.

### Summary of Updates

This version has been revised to reflect comments by referees during peer review. Major changes include: >Changing the assignment of postcranial remains from cf. Keuperodus brodiei to Elasmobranchii incertae sedis >Expanded justification of assignment of remains to K. brodiei >Inclusion of data on the holotype of K. brodiei >Expanded description of phylogenetic analysis In addition lots of smaller edits were made and additional references included.

